# Immunotherapeutic strategy to prevent progression and complications of acute rheumatic fever

**DOI:** 10.1101/2024.08.11.607519

**Authors:** Rukshan Ahamed Mohamed Rafeek, Natkunam Ketheesan, Michael F. Good, Manisha Pandey, Ailin Lepletier

**Affiliations:** School of Science & Technology, University of New England, New South Wales, Australia; Institute for Biomedicine and Glycomics, Griffith University, Gold Coast, Queensland, Australia

## Abstract

Acute rheumatic fever (ARF) is an autoimmune disease triggered by antibodies and T-cells targeting the Group A streptococcal (GAS, Strep A) bacterium, often leading to rheumatic heart disease (RHD). Long-term antibiotic therapy is recognized as a cornerstone of public health programs to prevent reinfection and progression of ARF. However, better tools to slow disease progression, and mitigate its lifelong consequences are required. Evidence obtained in a preclinical model suggests that this can be achieved. Using the rat autoimmune valvulitis model, we explored the potential of low-dose interleukin 2 (LD-IL-2) as an immunotherapeutic intervention. In this model injection of recombinant Strep A M5 protein (rM5) to Lewis rats induce autoimmune complications, cardiac tissue inflammation and conduction abnormalities. In animals injected with rM5 and treated with LD-IL-2, no cardiac functional or histological changes were observed. LD-IL-2 therapy effectively reduced the production of cross-reactive antibodies against cardiac tissue and induced a significant increase in classical regulatory T-cells (Treg) and CD8^+^ Tregs in the mediastinal (heart-draining) lymph nodes. These novel findings suggest LD-IL-2 will be an effective immunotherapeutic agent for treating ARF/RHD.

## Introduction

Rheumatic heart disease (RHD) is a neglected disease of poverty that affects 40 million people worldwide and leads to more than 350,000 deaths annually. It accounts for nearly 2% of all deaths from cardiovascular diseases and is the commonest cause of pediatric acquired heart disease (Watkins et al., 2017). In susceptible individuals, infections with Group A Streptococcus (GAS, Strep A) initiate an autoimmune process that leads to Acute Rheumatic Fever (ARF). Approximately 60% of patients who experience at least one episode of ARF will develop irreversible damage of the heart valves, which characterizes RHD (Good, 2020). Besides, up to 30% of patients with ARF develop Sydenham’s chorea (SC), a neurobehavioral condition characterized by involuntary choreiform movements and neuropsychiatric impairment. Although ARF is a systemic disease affecting multiple organs, only carditis results in lifelong consequences. As well as people from less economically developed countries indigenous populations in developed countries face significantly higher rates of diseases associated with Strep A infections. Among them, Indigenous Australians have some of the highest recorded incidences of ARF and RHD globally. Of the ARF diagnoses between 2016 and 2020, almost half (46%) were children aged 5-14 years. ARF rates in children aged under 15 years were generally higher among males, but in adults, ARF rates were generally higher among females (Welfare, 2022).

RHD is a sequela of group A streptococcal (Strep A) infections in which antibodies and T-cells raised against epitopes in the structural M-protein cross-react with host heart proteins resulting in heart damage. This process, known as molecular mimicry, is part of the host’s immune response to Strep A (Cunningham, 2019). Studies have identified cross-reactive antibodies and T helper cells - producers of IFN-ɣ and IL-17 (Th1 and Th17, respectively) – as key mediators of RHD (Cunningham, 2000; Sikder et al., 2018). For initiation of disease, evidence suggests that the initiation of disease may follow a two-hit hypothesis: (i) autoantibodies binding to and activating the valvular endothelium, and (ii) subsequent extravasation of T-cells through the activated endothelium into the valve. This leads to the recognition of autoantigens and amplification of the inflammatory process followed by tissue destruction (Carapetis et al., 2016). Deficiency of regulatory T-cells (Treg) was reported in patients with RHD as well as a high Th17/Treg ratio, a phenotype exacerbated in patients with multivalvular compared to univalvular involvement (Bas et al., 2014; Mukhopadhyay et al., 2013).

Currently, there is no specific treatment to arrest the progression of ARF to RHD. Additionally, there is no vaccine available to prevent repetitive Strep A infection, which can exacerbate ARF and drive the pathological process. For decades, patients diagnosed with ARF have been prescribed regular (4 weekly) and long-term (up to 10 years or more) administration of penicillin to prevent recurrent Strep A infections (Carapetis et al., 2016; Stollerman et al., 1955). Low dose interleukin-2 (LD-IL-2) therapy is known for its safety and efficacy in treating a number of autoimmune conditions, such as type 1 diabetes (Hartemann et al., 2013), systemic lupus erythematosus (He et al., 2016), and vasculitis associated with chronic hepatitis C virus. Unlike conventional immunosuppressive treatments, LD-IL-2 therapy promotes immune tolerance by specifically targeting Tregs without causing generalized immunosuppression that affects conventional T cells (Churlaud et al., 2014; Zhou et al., 2021). Five-day course of LD-IL-2 therapy with daily injections was shown to harness IL-2’s immunomodulatory properties and mitigate autoimmune responses in humans and in preclinical models (Churlaud et al., 2015; Grasshoff et al., 2021).

In this study, we utilize the Rat Autoimmune Valvulitis (RAV) model (Rafeek et al., 2022; Reynolds et al., 2023; Sikder et al., 2019) to investigate whether LD-IL-2 therapy can be repurposed to treat active ARF and prevent RHD.

## Results and Discussion

We have developed and characterized the RAV model to determine the early events that lead to the clinical syndromes of ARF complications. In this model, multiple injections of rats with a recombinant M protein from Strep A, type 5 (rM5, Strep A M5 is a classical “rheumatogenic” strain) emulsified in Freund’s adjuvant induces antibodies and T-cells that cross-react with cardiac tissue (Gorton et al., 2009; Lymbury et al., 2003; Sikder et al., 2019). The histological, immunological, and functional changes in the hearts resemble that of RHD, including Aschoff’s nodules. To investigate whether LD-IL-2 therapy can be repurposed to treat carditis, we compared PBS-injected control rats with rats injected with rM5, which were either treated with LD-IL-2 or left untreated. Treatment regimen consisted of repeated subcutaneous injections of LD-IL-2 after rats have received the first (Day 8) or last (Day 21) booster with Strep A rM5 (Fig. 1A). For both regimens, LD-IL-2 was administered daily for 5 days, followed by 4 injections with a 2-day interval between each injection. Euthanasia was performed on Day 35, either 11 days following the conclusion of LD-IL-2 treatment (Day 8) or 2 days following the end of treatment (Day 21).

**Figure 1.**
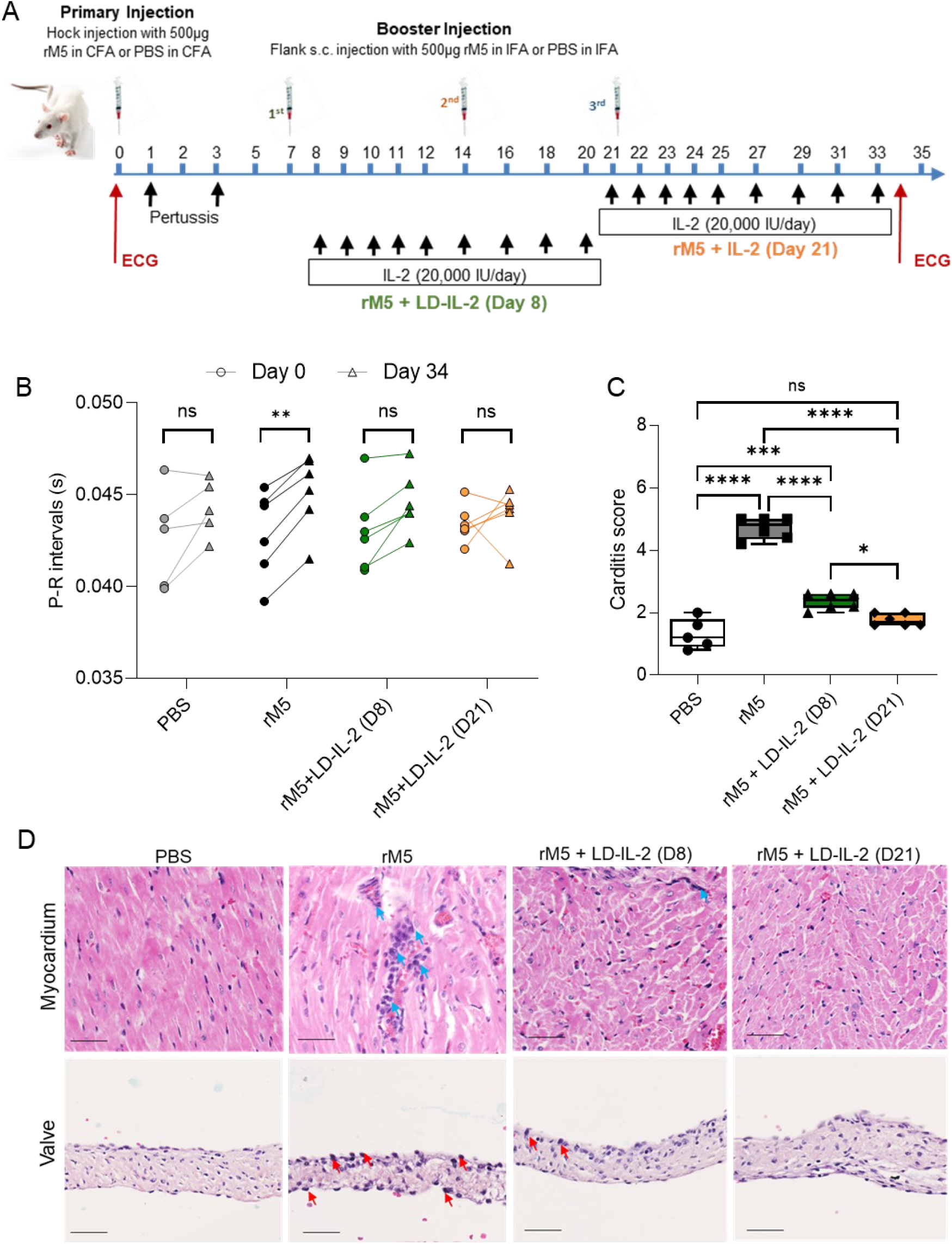
LD-IL-2 therapy induces functional improvement of the heart and reduces carditis. (A) Experimental timeline summarizing development of low-dose IL-2 (LD-IL-2) therapeutics for ARF/RHD using the Rat Autoimmune Valvulitis (RAV) model. Rats received 3 boosters with Strep A rM5 at weekly intervals. LD-IL-2 treatment began either on Day 8 or Day 21 post-primary injection with Strep A rM5. Rats were euthanized at Day 35 for assessment of carditis and immune responses. (B) Functional assessment of the heart before (•) and after (Δ) injection of PBS or rM5 by ECG. Antigen-injected rats (n=5-6 females/grp) received LD-IL-2 therapy at Day 8 (D8) or Day 21 (D21) post-primary injection or were left untreated. Statistical analysis performed by two-way ANOVA. n.s. non-significant, ** p<0.01. (C) Inflammatory changes in the myocardium and valvular tissue, characterized by mononuclear cell infiltration, were scored in rats treated with PBS, rM5, and LD-IL-2 therapy started at Day 8 or Day 21 post-primary rM5 injection. Box-and-whisker plot show min to max carditis scores within each treatment group. Statistical analysis for carditis score were obtained by one-way ANOVA. n.s. non-significant, *p<0.05, *** p<0.005, ***p<0.001. (D) Representative histological images of myocardium and valvular tissues. Arrows indicate inflammatory cell infiltration in the myocardium (blue arrows) and valves (red arrows). Scale bar = 50µm.

### LD-IL-2 therapy prevents the functional and histological cardiac changes associated with RHD

An important characteristic of the RAV model is its ability to induce observable functional and histological changes to the heart, mirroring those clinically observed in patients with ARF/RHD. To measure conduction abnormalities and assess cardiac function, all rats underwent electrocardiography (ECG) before being culled on Day 35. Baseline heart function prior to the first Strep A rM5 injection (Day 0) was used to compare functional changes. Peak values of P and R points at three different segments of ECG from each rat were individually measured and analyzed. A prolongation of the P-R interval was found in all rM5-treated rats, indicating a conduction abnormality in the heart (Fig. 1B). In contrast, following both LD-IL-2 therapeutic regimens, rats demonstrated no change to P-R interval.

To investigate the effect of LD-IL-2 therapy on inflammatory changes associated with ARF/RHD, infiltrating mononuclear cells were assessed by hematoxylin and eosin (H&E) staining. The extent of inflammation in the myocardium and valves was expressed as a “carditis score” which was based on the number of mononuclear cell infiltrates and focal lesions in both the myocardium and mitral valve. Carditis scoring was undertaken using a scoring matrix previously developed (Reynolds et al., 2023). An increased mononuclear cell infiltration was observed in rats injected with rM5 compared to PBS control group. In contrast, minimal mononuclear cell infiltration was observed in rM5 injected rats treated with LD-IL-2 from either Day 8 or Day 21. Both LD-IL-2 treatment regimens resulted in a significant reduction in carditis (Fig. 1C and 1D). Although the carditis scores were statistically different between rats treated with the Day8 regimen and Day 21 regimen, the absolute differences were small.

It is possible that the timing of LD-IL-2 treatment in ARF may influence the effectiveness of LD-IL-2 therapy, depending on the stage and chronicity of the autoimmune process. Notwithstanding this, the present study demonstrates that commencing IL-2 immunotherapy either early or later in the time course of M5 protein exposure and hence disease onset, were both effective in treating any existing disease.

### Reduction of cross-reactive M-protein antibodies following LD-IL-2 therapy

Tissue cross-reactive antibodies against Strep A antigens play a significant role in initiating the autoimmune pathology in ARF/RHD. A key aspect of the RAV model is its ability to generate antibodies that cross-react with cardiac and neuronal tissue. To investigate whether LD-IL-2 therapy can inhibit the induction of cross-reactive antibodies, we conducted enzyme-linked immunosorbent assay (ELISA) against cardiac myosin and tropomyosin (heart muscle), laminin (heart valve extracellular matrix), and dopamine receptors type 1 and 2, tubulin and lysoganglioside (brain). Although the rM5-injected group exhibited significant cross-reactivity with all tissue proteins compared to the negative control PBS group, no cross-reactivity to cardiac tissues was observed for the antisera from both LD-IL-2 treatment groups (Fig. 2). LD-IL-2 also resulted in significant reduction in the levels of IgG recognizing connective and neuronal proteins. No significant difference was observed between both treatment groups, and treatment started at Day 21 resulted in antibody levels similar to the control group (Fig. 2). In ARF/RHD, tissue cross-reactive antibodies activate the valvular endothelium and trigger carditis. In SC, cross reactive antibodies that transverse the compromised blood brain barrier and bind to neuronal proteins in the basal ganglia are considered to be the cause for the onset of neurobehavioral symptoms (Carapetis et al., 2016; Soller et al., 2023). Autoantibodies against dopamine receptors may lead to a receptor imbalance and induce greater sensitivity to dopamine signaling potentially leading to neuropsychiatric symptoms in SC (Ben-Pazi et al., 2013).

**Figure 2.**
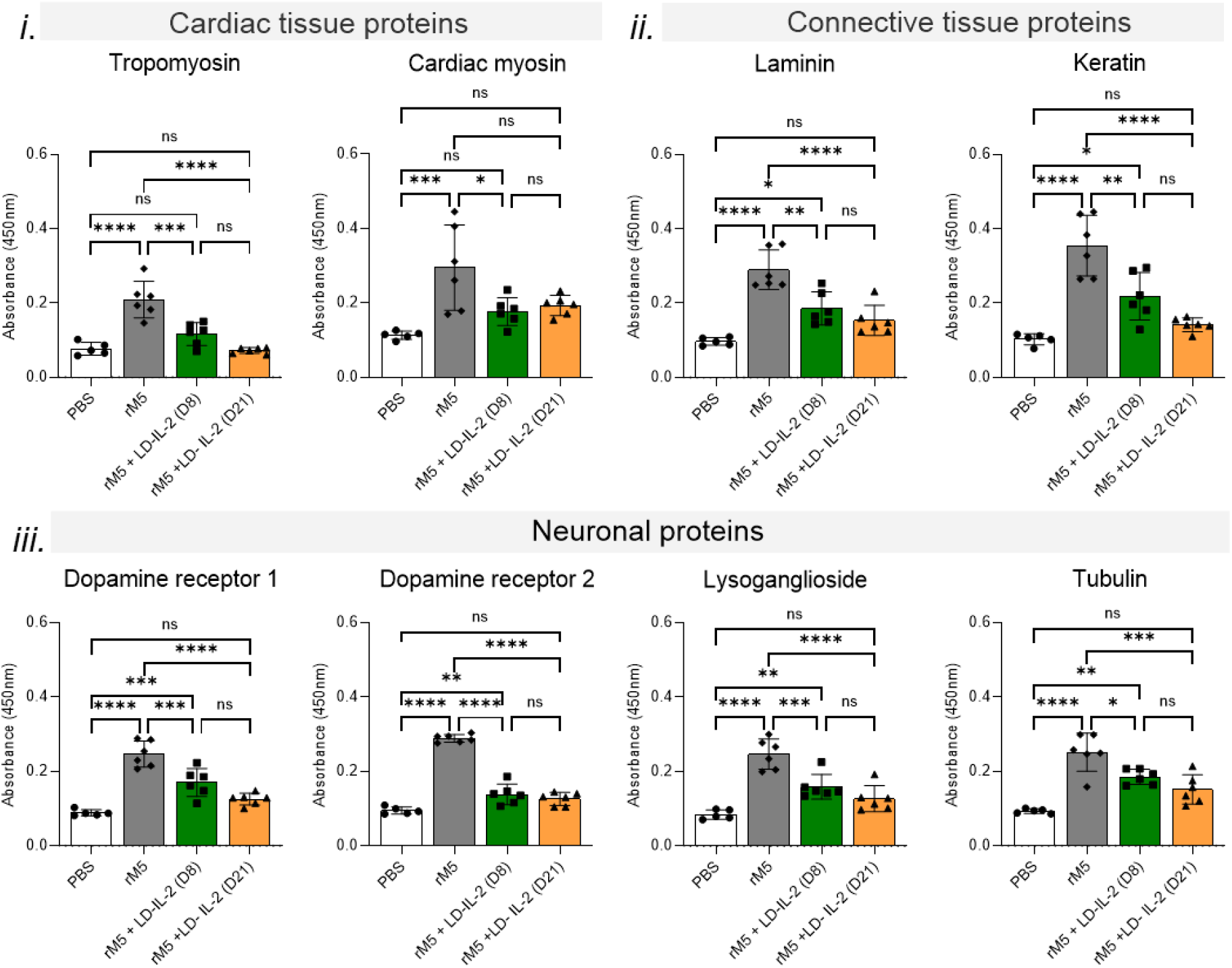
LD-IL-2 therapy reduces serum IgG cross-reactivity to host cardiac and neuronal tissue proteins. Serum IgG levels against *i*. cardiac proteins (tropomyosin and cardiac myosin), ii. connective tissues (laminin and keratin) and *iii*. neuronal proteins (dopamine receptors 1 and 2, lysoganglioside and tubulin). Absorbance values of Day-35 rat sera at 1:400 dilution is shown (n=5-6 females/grp). Statistical analysis performed by one-way ANOVA with Dunnett multiple comparison test. n.s. non-significant, *p < 0.05, **p < 0.01, ***p<0.005, ****p< 0.001. Bars represent standard deviation (SD).

LD-IL-2 can indirectly affect B cell differentiation and antibody production. By promoting a regulatory environment through Tregs, LD-IL-2 helps to control the differentiation of B cells into antibody-secreting plasma cells that are enriched for a regulatory B cell signature (Inaba et al., 2023), thereby reducing the production of self-reactive antibodies. Besides, LD-IL-2 suppress the activation and differentiation of T follicular helper (Tfh) cells (Ballesteros-Tato, 2014; Liang et al., 2021). This modulation helps to limit excessive germinal center responses that contribute to autoantibody production and promotes immune tolerance, highlighting LD-IL-2 as a potential therapeutic strategy for managing ARF/RHD.

### LD-IL-2 therapy selectively expands regulatory T-cells (Treg) in mediastinal lymph nodes

Tregs are a specialized T-cell subset that regulates the immune system, maintaining homeostasis and self-tolerance (Singh et al., 2023; Wong et al., 2021). Decreased numbers or defective function of Tregs has been implicated in the pathogenesis of various autoimmune diseases (Dominguez-Villar and Hafler, 2018; Weerakoon et al., 2021). Although higher doses of IL-2 therapy have been traditionally approved for cancer therapy, the concept of using LD-IL-2 for treating autoimmune and inflammatory diseases emerged due to its potential to induce Treg cells compared to conventional T-cells and NK cells (Malek and Castro, 2010).

We investigated the impact of LD-IL-2 therapy on Treg and conventional T-cells from mediastinal (heart-draining) lymph nodes using flow cytometry. Striking, LD-IL-2 therapy resulted in 50% increase of classical Treg (CD4^+^CD25^+^FoxP3^+^) in lymph nodes of rats injected with rM5 (Fig. 3A). This is consistent with the expansion achieved in other rodent models assessing efficacy of LD-IL-2 therapy for treating autoimmune disorders (Zhou et al., 2021). Noteworthy, an increase of rare CD8^+^ Tregs (CD8^+^CD25^+^FoxP3^+^) was observed when rats were treated with LD-IL-2 from Day 21, but not when they were treated from Day 8 (Fig. 3B). In contrast to classical Tregs, the origin and roles of CD8^+^ Treg in the pathogenesis of autoimmune diseases are less understood. While naturally occurring CD8^+^Tregs are rare, they can be induced *in vivo* (Flippe et al., 2019) and are generally considered to maintain less stable expression of FoxP3 than classical Tregs (Iamsawat et al., 2018; Wang et al., 2021). Nonetheless, CD8^+^ Tregs show heightened sensitivity to IL-2 for proliferation compared to T effector cells and have been induced by LD-IL-2 therapy in both mice and humans, leading to benefits in alleviating autoimmune disorders (Aoyama et al., 2012; Churlaud et al., 2015; Dinesh et al., 2010). Similar to their CD4^+^ counterpart, CD8^+^ Tregs have been shown to play important roles in immune regulation suppressing immune responses through a variety of mechanisms that include secretion of cytokines, cell-to-cell contact, induction of a tolerogenic phenotype in APCs that can then induce classical Tregs, and cytotoxic activity (Mishra et al., 2021; Vieyra-Lobato et al., 2018).

**Figure 3.**
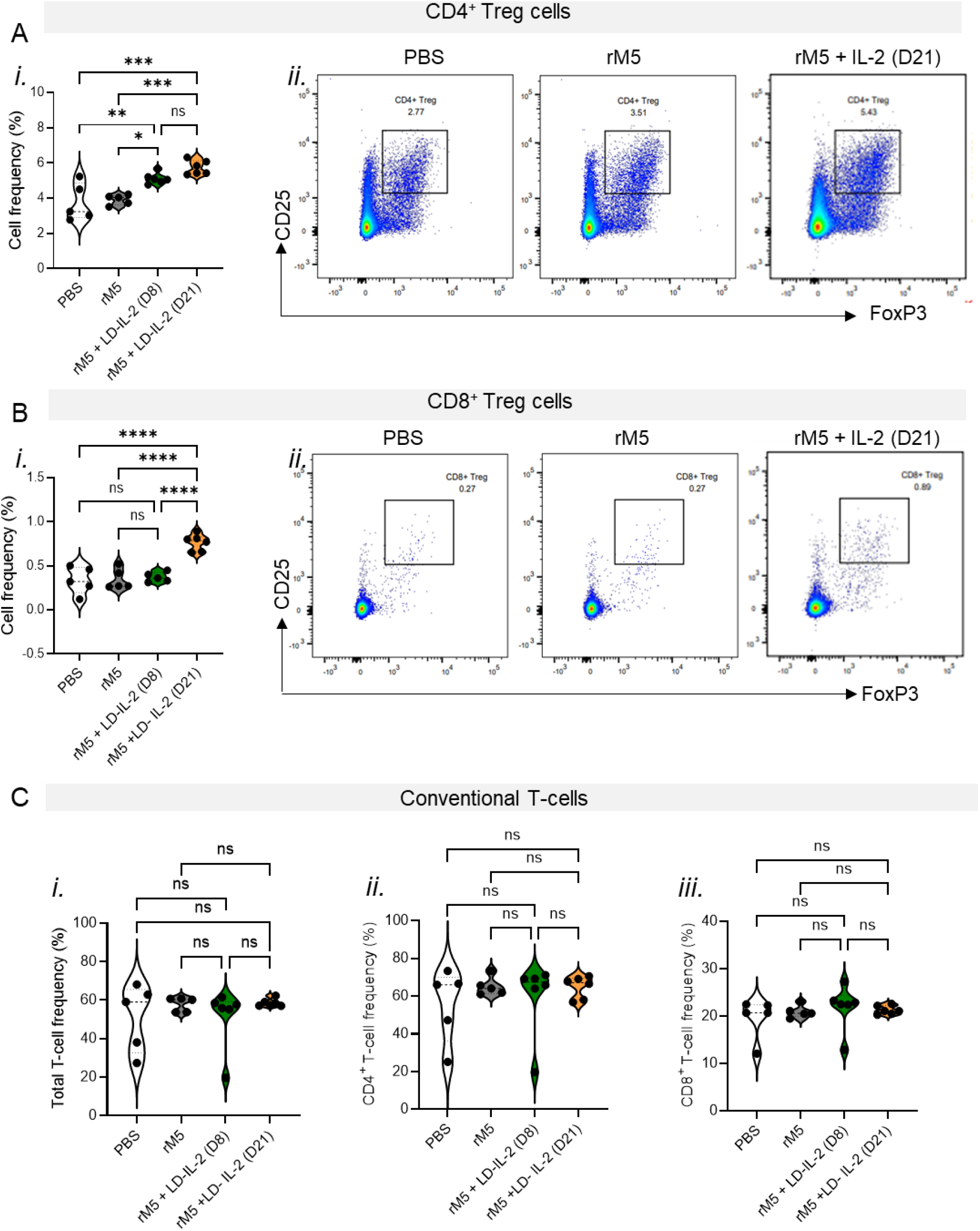
LD-IL-2 therapeutic efficacy is associated with a targeted increase in Treg cells. Flow cytometry analysis of T cells in mediastinal lymph nodes. (A) *i*. Violin plot is used to compare the proportion on CD4^+^ Treg (CD3^+^CD4^+^CD25^+^FoxP3^+^) between rats treated with PBS, rM5 only and rM5+LD-IL-2 (n=5-6). Each symbol represents an individual rat analyzed. *ii*. Representative gating strategy for defining CD4^+^ Treg. (B) *i*. Proportion on CD8+ Treg (CD3^+^CD8^+^CD25^+^FoxP3^+^) between treatment groups. Each symbol represents an individual rat analyzed (n=5-6 females/grp). *ii*. Representative gating strategy for defining CD8^+^ Treg (C) Violin plot is used to compare the proportion of *i*. total T-cells, *ii*. CD4^+^ T-cells and *iii*. CD8^+^ T-cells between rats treated with PBS, rM5 only and rM5+LD-IL-2. Each symbol represents an individual donor analyzed (n=5-6 females/grp). Non-parametric one-way ANOVA test with Bonferroni correction. ns = non-significant, *P < 0.05, **P < 0.01, ***P < 0.005, ****p< 0.001. Non-parametric one-way ANOVA test with Bonferroni correction.

In our study, we did not observe induction of CD8^+^ Tregs when rats were treated with LD-IL-2 therapy starting from Day 8. This lack of induction is likely due to the instability of FoxP3 expression in induced Tregs, which could not be sustained until the experiment endpoint, at Day 35. Although administrating LD-IL-2 expanded Tregs and reduced disease severity in a several autoimmune diseases (Hartemann et al., 2013; He et al., 2016; Zheng et al., 2022), IL-2 is also a growth factor for potentially pathogenic conventional T-cells due to the widespread expression of IL-2R subunits. Thus, IL-2’s dual role in promoting both tolerance and activation makes predicting its therapeutic effects challenging. Moreover, low dose IL-2 has been demonstrated to exhibit limited or no therapeutic efficacy in alloimmune conditions (Lim et al., 2023) as well as heterogeneity in clinical responsiveness in various autoimmune diseases (Grasshoff et al., 2021). Furthermore, a therapeutic role for low dose IL-2 in autoimmune diseases mediated by bacterial antigens and subsequent molecular mimicry has not been previously described.

To determine if the LD-IL-2 therapeutic regimen used in this study could also expand conventional T-cells, we analyzed the frequencies of total conventional T-cells and the CD4^+^ and CD8^+^ T-cell subsets. Notably, LD-IL-2 therapy did not alter the frequencies of conventional T-cells but specifically expanded Tregs (Fig. 3C) suggesting that LD-IL-2 is a safe and specific approach for treating ARF and preventing RHD.

Treg cells constitutively express high levels of the heterotrimeric high-affinity IL-2 receptor (IL-2R) complex and the highest expression of the high-affinity α chain (CD25) of the IL-2R complex, making them particularly sensitive to even small amounts of IL-2 in the body. However, a genetic deficiency in the IL-2/IL-2R pathway may leads to systemic autoimmunity (Sadlack et al., 1993). Due to this unique property LD-IL-2 therapy has become an emerging immunotherapeutic approach for treating certain autoimmune conditions. In this study we experimentally demonstrated that repeated injection of LD-IL-2 following rM5 injection resulted in expansion of both CD4^+^ and CD8^+^ Tregs and thereby reduced the cardiac inflammation. The short half-life of IL-2 necessitates a course of repetitive injections to maintain therapeutic levels of Tregs for treatment of autoimmune diseases (Lotze et al., 1985). The findings of this study suggest that LD-IL-2 therapy holds promise for managing ARF by effectively reducing autoimmune responses and potentially preventing carditis. By targeting immune system dysregulation, LD-IL-2 could represent a valuable therapeutic approach. This approach may lead to improved clinical outcomes and reduced incidence of cardiac complications in individuals affected by ARF. This approach may lead to improved clinical outcomes and reduced incidence of cardiac complications in individuals affected by ARF.

## Material and methods

### Animals

All experimental protocols involving animals were approved by the Animal Ethics Committee of University of New England (UNE) (ARA23-009). Female Lewis rats (LEW/SsN; Albino:a,h,c:RT^1^) aged 4 to 6 weeks were purchased from the Centre for Animal Research and Teaching at UNE and acclimatised for 5 days prior to experiments.

### Preparation of recombinant Strep A M5 proteins

Recombinant M5 protein of Strep A (rM5) was cloned and purified as described previously (Gorton. et al., 2009). Briefly, Strep A *emm5* was cloned into pREP4 vector and expressed in *E*.*coli* BL21. Recombinant M proteins were purified using Ni-NTA resin. Lipopolysaccharide (LPS) contamination in recombinant protein preparations was removed by Triton X-114 assisted LPS extraction as previously described (Teodorowicz et al., 2017). The rM5 protein preparations was determined to be free of LPS using Pierce Chromogenic Endotoxin Quantification Kit (ThermoFisher, USA) as per manufacturer’s instructions.

### Induction of autoimmune carditis and valvulitis

For induction of carditis, rats were injected with 0.5 mg/100 μL of recombinant Strep A M5 protein emulsified in Complete Freund’s Adjuvant (CFA) or PBS emulsified in CFA. Baseline Electrocardiography (ECG) was performed prior to injection. All priming injections were performed under isoflurane inhalation anaesthesia (5% induction and 2% maintenance) in the hock as described previously (Gorton et al., 2010). On day 1 and 3 all rats were intraperitoneally (i.p.) injected with 300 ng of *Bordetella pertussis* (BPTx) toxin (Gibco, USA) in 200 μL PBS. Rats were boosted with respective antigens in Incomplete Freund’s Adjuvants (IFA) on the flank on days 7, 14, and 21. The rats were euthanized on day 35 from the day of primary injection with the overdose of sodium pentobarbital (260mg/kg, i.p), blood was collected by cardiac puncture and heart tissue was collected and fixed with 4% paraformaldehyde for histopathological analysis. Heart draining mediastinal lymph nodes were collected for flow cytometry analysis.

### Low dose interleukin-2 (LD-IL-2) treatment

Twenty-three age matched female Lewis rats were divided into four treatment groups (n=5/6); (i) PBS, (ii) Strep A rM5, (iii) Strep A rM5 + IL-2 (D8) and (iv) Strep A rM5 + IL-2 (D21). Recombinant Human IL-2 (#200-02, Peprotech, USA) was reconstituted in 100mM acetic acid and further diluted in PBS containing 0.1% Bovine Serum Albumin (BSA) to achieve 20,000 IU concentration. Rats in group iii received s.c injection of LD-IL-2 from Day 8 of primary injection to Day 24 on every other day. Rats in group iv received s.c injection of LD-IL-2 from Day 21 of primary injection to Day 34 on every other day (Supplementary Figure 1).

### Electrocardiography (ECG)

ECG traces were recorded for 1-2 min using Bio Amp with PowerLab data acquisition system (ADInstruments, USA) in all rats prior to injection and a day before euthanasia to assess the conduction abnormalities in the heart. Peak values of P and R points at three different segments of ECG from each rat was individually extracted and analysed (Rafeek et al., 2021).

### Histopathological analysis of cardiac tissue

To examine the extent of inflammation, formalin-fixed paraffin-embedded (FFPE) cardiac tissue was processed, embedded in paraffin, sectioned, and stained with Harris H&E using standard procedures as previously described (Gorton. et al., 2009). Slides were examined microscopically for infiltration of inflammatory cells as evidence of myocarditis or valvulitis. The extent of inflammation was expressed as a “carditis score” based on the number of inflammatory cells and focal lesions from each rat (Reynolds et al., 2023). The scoring system assesses and scores valvular and myocardial sections (5 randomly selected areas per animal) from the different groups. These sections were blindly scored using a validated semi-quantitative scoring system.

### Detection of tissue-reactive serum antibodies

Serum IgG antibody against purified host proteins including cardiac myosin, tropomyosin, laminin, keratin, dopamine receptors 1&2, lysoganglioside_GM1_ and tubulin (Sigma, USA) were determined using indirect ELISA as described previously (Rafeek et al., 2022). Briefly purified proteins were coated onto Maxisorp 96-well plates (Nunc, USA) in carbonate-bicarbonate buffer. Plates were blocked and incubated with individual rat sera in duplicate at 1:400 dilution. Following repeated washing of the plates, each well was incubated with 100 µL of goat anti-rat IgG HRP conjugated secondary antibody (1:5000; Jackson Immunoresearch, USA) for an hour at room temperature. Each well was washed and incubated for 20 min with 100 µL of SIGMAFAST™ OPD (Sigma, USA). Absorbance was measured at 450 nm using SpectraMax M2/M2e (Molecular Devices, USA).

### Multiparametric flow cytometry for analysis of circulating Treg

Mediastinal lymph nodes were dissociated into single-cells suspension were obtained from mediastinal lymph nodes using a plunger from a sterile syringe. Red blood cells were lysed with ACK lysis buffer and washed with RPMI (Thermo Fisher Scientific, AUS). The cell populations were determined by flow cytometry. The cells were surface-stained with a master mix containing dead cell exclusion dye (Zombie violet), CD4-APC-Cy7, CD3-PerCP, CD8-PE-Cy7 and CD25-APC. Cells were stained on ice in the dark for 40 min. Intranuclear staining for FoxP3-PE was achieved using FoxP3 Fix/Perm Buffer Set for nuclear staining (BioLegend, San Diego, California, USA). Samples were washed in PBS and acquired Samples were acquired by a BD Fortessa multiparametric flow cytometer (BD Biosciences, Franklin Lakes, New Jersey, USA). Data were analyzed using FlowJo V.10.7 (BD).

### Statistical analysis

Descriptive statistics of results were made using GraphPad Prism version 8.0. Antibody titre data are presented as geometric mean. One-way ANOVA with Dunnett post hoc method for multiple comparisons was used for pairwise comparisons. Results were taken as significant with P-values <0.05 or <0.01. Two-way ANOVA was used to assess ECG differences between rats. All data with p-values <0.05 were considered significant.

## Acknowledgements

The authors acknowledge Dr Rhonda Davey for the assistance with histology and Dr Daniel Ebert for logistical support and Mr Mohamed Raguib Munif for the assistance with animal experiments. Rukshan AM Rafeek is supported by an NHMRC Ideas Grant (APP 2010336) to Natkunam Ketheesan. AL is supported by an NHMRC project grant (APP1160379) awarded to Manisha Pandey. The work described in this manuscript was supported by Griffith University Internal Grant to Ailin Lepletier.

## Author contributions

A.L.and M.P. equally contributed to this work. A.L, R.A.M.R, M.P, N.K and M.F.G conceptualised and designed the experiments. R.A.M.R and A.L performed experiments and analysed data. All authors contributed to the writing of the manuscript and have read and approved the submitted version.

## References

Aoyama, A., D. Klarin, Y. Yamada, S. Boskovic, O. Nadazdin, K. Kawai, D. Schoenfeld, J.C. Madsen, A.B. Cosimi, G. Benichou, and T. Kawai. 2012. Low-dose IL-2 for In vivo expansion of CD4+ and CD8+ regulatory T cells in nonhuman primates. Am J Transplant 12:2532–2537.

Ballesteros-Tato, A. 2014. Beyond regulatory T cells: the potential role for IL-2 to deplete T-follicular helper cells and treat autoimmune diseases. Immunotherapy 6:1207–1220.

Bas, H.D., K. Baser, E. Yavuz, H.A. Bolayir, B. Yaman, S. Unlu, A. Cengel, E.U. Bagriacik, and R. Yalcin. 2014. A shift in the balance of regulatory T and T helper 17 cells in rheumatic heart disease. J Investig Med 62:78–83.

Ben-Pazi, H., J.A. Stoner, and M.W. Cunningham. 2013. Dopamine receptor autoantibodies correlate with symptoms in Sydenham’s chorea. PLoS One 8:e73516.

Carapetis, J.R., A. Beaton, M.W. Cunningham, L. Guilherme, G. Karthikeyan, B.M. Mayosi, C. Sable, A. Steer, N. Wilson, R. Wyber, and L. Zuhlke. 2016. Acute rheumatic fever and rheumatic heart disease. Nat Rev Dis Primers 2:15084.

Churlaud, G., V. Jimenez, J. Ruberte, M. Amadoudji Zin, G. Fourcade, G. Gottrand, E. Casana, B. Lambrecht, B. Bellier, E. Piaggio, F. Bosch, and D. Klatzmann. 2014. Sustained stimulation and expansion of Tregs by IL2 control autoimmunity without impairing immune responses to infection, vaccination and cancer. Clin Immunol 151:114–126.

Churlaud, G., F. Pitoiset, F. Jebbawi, R. Lorenzon, B. Bellier, M. Rosenzwajg, and D. Klatzmann. 2015. Human and Mouse CD8(+)CD25(+)FOXP3(+) Regulatory T Cells at Steady State and during Interleukin-2 Therapy. Front Immunol 6:171.

Cunningham, M.W. 2000. Pathogenesis of group A streptococcal infections. Clin Microbiol Rev 13:470–511.

Cunningham, M.W. 2019. Molecular Mimicry, Autoimmunity, and Infection: The Cross-Reactive Antigens of Group A Streptococci and their Sequelae. Microbiol Spectr 7:

Dinesh, R.K., B.J. Skaggs, A. La Cava, B.H. Hahn, and R.P. Singh. 2010. CD8+ Tregs in lupus, autoimmunity, and beyond. Autoimmun Rev 9:560–568.

Flippe, L., S. Bezie, I. Anegon, and C. Guillonneau. 2019. Future prospects for CD8(+) regulatory T cells in immune tolerance. Immunol Rev 292:209–224.

Good, M.F. 2020. Streptococcus: An organism causing diseases beyond neglect. PLoS Negl Trop Dis 14:e0008095.

Gorton, D., S. Blyth, J.G. Gorton, B. Govan, and N. Ketheesan. 2010. An alternative technique for the induction of autoimmune valvulitis in a rat model of rheumatic heart disease. J Immunol Methods 355:80–85.

Gorton, D., B. Govan, C. Olive, and N. Ketheesan. 2009. B- and T-cell responses in group a streptococcus M-protein-or Peptide-induced experimental carditis. Infect Immun 77:2177–2183.

Gorton., B. Govan, C. Olive, and N. Ketheesan. 2009. B- and T-cell responses in group a streptococcus M-protein-or Peptide-induced experimental carditis. Infect Immun 77:2177–2183.

Grasshoff, H., S. Comduhr, L.R. Monne, A. Muller, P. Lamprecht, G. Riemekasten, and J.Y. Humrich. 2021. Low-Dose IL-2 Therapy in Autoimmune and Rheumatic Diseases. Front Immunol 12:648408.

Hartemann, A., G. Bensimon, C.A. Payan, S. Jacqueminet, O. Bourron, N. Nicolas, M. Fonfrede, M. Rosenzwajg, C. Bernard, and D. Klatzmann. 2013. Low-dose interleukin 2 in patients with type 1 diabetes: a phase 1/2 randomised, double-blind, placebo-controlled trial. Lancet Diabetes Endocrinol 1:295–305.

He, J., X. Zhang, Y. Wei, X. Sun, Y. Chen, J. Deng, Y. Jin, Y. Gan, X. Hu, R. Jia, C. Xu, Z. Hou, Y.A. Leong, L. Zhu, J. Feng, Y. An, Y. Jia, C. Li, X. Liu, H. Ye, L. Ren, R. Li, H. Yao, Y. Li, S. Chen, X. Zhang, Y. Su, J. Guo, N. Shen, E.F. Morand, D. Yu, and Z. Li. 2016. Low-dose interleukin-2 treatment selectively modulates CD4(+) T cell subsets in patients with systemic lupus erythematosus. Nat Med 22:991–993.

Iamsawat, S., A. Daenthanasanmak, J.H. Voss, H. Nguyen, D. Bastian, C. Liu, and X.Z. Yu. 2018. Stabilization of Foxp3 by Targeting JAK2 Enhances Efficacy of CD8 Induced Regulatory T Cells in the Prevention of Graft-versus-Host Disease. J Immunol 201:2812–2823.

Inaba, A., Z.K. Tuong, T.X. Zhao, A.P. Stewart, R. Mathews, L. Truman, R. Sriranjan, J. Kennet, K. Saeb-Parsy, L. Wicker, F. Waldron-Lynch, J. Cheriyan, J.A. Todd, Z. Mallat, and M.R. Clatworthy. 2023. Low-dose IL-2 enhances the generation of IL-10-producing immunoregulatory B cells. Nat Commun 14:2071.

Liang, K., J. He, Y. Wei, Q. Zeng, D. Gong, J. Qin, H. Ding, Z. Chen, P. Zhou, P. Niu, Q. Chen, C. Ding, L. Lu, X.X. Chen, Z. Li, N. Shen, D. Yu, and J. Deng. 2021. Sustained low-dose interleukin-2 therapy alleviates pathogenic humoral immunity via elevating the Tfr/Tfh ratio in lupus. Clin Transl Immunology 10:e1293.

Lim, T.Y., E. Perpinan, M.C. Londono, R. Miquel, P. Ruiz, A.S. Kurt, E. Kodela, A.R. Cross, C. Berlin, J. Hester, F. Issa, A. Douiri, F.H. Volmer, R. Taubert, E. Williams, A.J. Demetris, A. Lesniak, G. Bensimon, J.J. Lozano, M. Martinez-Llordella, T. Tree, and A. Sanchez-Fueyo. 2023. Low dose interleukin-2 selectively expands circulating regulatory T cells but fails to promote liver allograft tolerance in humans. J Hepatol 78:153–164.

Lymbury, R.S., C. Olive, K.A. Powell, M.F. Good, R.G. Hirst, J.T. LaBrooy, and N. Ketheesan. 2003. Induction of autoimmune valvulitis in Lewis rats following immunization with peptides from the conserved region of group A streptococcal M protein. J Autoimmun 20:211–217.

Malek, T.R., and I. Castro. 2010. Interleukin-2 receptor signaling: at the interface between tolerance and immunity. Immunity 33:153–165.

Mishra, S., W. Liao, Y. Liu, M. Yang, C. Ma, H. Wu, M. Zhao, X. Zhang, Y. Qiu, Q. Lu, and N. Zhang. 2021. TGF-beta and Eomes control the homeostasis of CD8+ regulatory T cells. J Exp Med 218:

Mukhopadhyay, S., S. Varma, S. Gade, J. Yusuf, V. Trehan, and S. Tyagi. 2013. Regulatory T-cell deficiency in rheumatic heart disease: a preliminary observational study. J Heart Valve Dis 22:118–125.

Rafeek, R.A., A.S. Hamlin, N.M. Andronicos, C.S. Lawlor, D.J. McMillan, K.S. Sriprakash, and N. Ketheesan. 2022. Characterization of an experimental model to determine streptococcal M protein-induced autoimmune cardiac and neurobehavioral abnormalities. Immunol Cell Biol 100:653–666.

Rafeek, R.A.M., C.M. Lobbe, E.C. Wilkinson, A.S. Hamlin, N.M. Andronicos, D.J. McMillan, K.S. Sriprakash, and N. Ketheesan. 2021. Group A streptococcal antigen exposed rat model to investigate neurobehavioral and cardiac complications associated with post-streptococcal autoimmune sequelae. Animal Model Exp Med 4:151–161.

Reynolds, S., R.A.M. Rafeek, A. Hamlin, A. Lepletier, M. Pandey, N. Ketheesan, and M.F. Good. 2023. Streptococcus pyogenes vaccine candidates do not induce autoimmune responses in a rheumatic heart disease model. NPJ Vaccines 8:9.

Sikder, S., G. Price, M.A. Alim, A. Gautam, R. Scott Simpson, C. Margaret Rush, B. Lee Govan, and N. Ketheesan. 2019. Group A streptococcal M-protein specific antibodies and T-cells drive the pathology observed in the rat autoimmune valvulitis model. Autoimmunity 52:78–87.

Sikder, S., N.L. Williams, A.E. Sorenson, M.A. Alim, M.E. Vidgen, N.J. Moreland, C.M. Rush, R.S. Simpson, B.L. Govan, R.E. Norton, M.W. Cunningham, D.J. McMillan, K.S. Sriprakash, and N. Ketheesan. 2018. Group G Streptococcus Induces an Autoimmune Carditis Mediated by Interleukin 17A and Interferon gamma in the Lewis Rat Model of Rheumatic Heart Disease. J Infect Dis 218:324–335.

Singh, T.P., C. Farias Amorim, V.M. Lovins, C.W. Bradley, L.P. Carvalho, E.M. Carvalho, E.A. Grice, and P. Scott. 2023. Regulatory T cells control Staphylococcus aureus and disease severity of cutaneous leishmaniasis. J Exp Med 220:

Soller, T., K.V. Roberts, B.F. Middleton, and A.P. Ralph. 2023. Sydenham chorea in the top end of Australia’s Northern Territory: A 20-year retrospective case series. J Paediatr Child Health 59:1210–1216.

Stollerman, G.H., J.H. Rusoff, and I. Hirschfeld. 1955. Prophylaxis against group A streptococci in rheumatic fever; the use of single monthly injections of benzathine penicillin G. N Engl J Med 252:787–792.

Teodorowicz, M., O. Perdijk, I. Verhoek, C. Govers, H.F. Savelkoul, Y. Tang, H. Wichers, and K. Broersen. 2017. Optimized Triton X-114 assisted lipopolysaccharide (LPS) removal method reveals the immunomodulatory effect of food proteins. PLoS One 12:e0173778.

Vieyra-Lobato, M.R., J. Vela-Ojeda, L. Montiel-Cervantes, R. Lopez-Santiago, and M.C. Moreno-Lafont. 2018. Description of CD8(+) Regulatory T Lymphocytes and Their Specific Intervention in Graft-versus-Host and Infectious Diseases, Autoimmunity, and Cancer. J Immunol Res 2018:3758713.

Wang, W., T. Hong, X. Wang, R. Wang, Y. Du, Q. Gao, S. Yang, and X. Zhang. 2021. Newly Found Peacekeeper: Potential of CD8+ Tregs for Graft-Versus-Host Disease. Front Immunol 12:764786.

Watkins, D.A., C.O. Johnson, S.M. Colquhoun, G. Karthikeyan, A. Beaton, G. Bukhman, M.H. Forouzanfar, C.T. Longenecker, B.M. Mayosi, G.A. Mensah, B.R. Nascimento, A.L.P. Ribeiro, C.A. Sable, A.C. Steer, M. Naghavi, A.H. Mokdad, C.J.L. Murray, T. Vos, J.R. Carapetis, and G.A. Roth. 2017. Global, Regional, and National Burden of Rheumatic Heart Disease, 1990-2015. N Engl J Med 377:713–722.

Welfare, A.I.o.H.a. 2022. Acute rheumatic fever and rheumatic heart disease in Australia 2016–2020. Australian Government catalogue number CVD 95, AIHW,:

Wong, H.S., K. Park, A. Gola, A.P. Baptista, C.H. Miller, D. Deep, M. Lou, L.F. Boyd, A.Y. Rudensky, P.A. Savage, G. Altan-Bonnet, J.S. Tsang, and R.N. Germain. 2021. A local regulatory T cell feedback circuit maintains immune homeostasis by pruning self-activated T cells. Cell 184:3981–3997 e3922.

Zheng, X., R. Su, F. Hu, Y. Liu, X. Li, C. Gao, and C. Wang. 2022. Low-dose IL-2 therapy restores imbalance between Th17 and regulatory T cells in patients with the dermatomyositis combined with EBV/CMV viremia. Autoimmun Rev 21:103186.

Zhou, P., J. Chen, J. He, T. Zheng, J. Yunis, V. Makota, Y.O. Alexandre, F. Gong, X. Zhang, W. Xie, Y. Li, M. Shao, Y. Zhu, J.E. Sinclair, M. Miao, Y. Chen, K.R. Short, S.N. Mueller, X. Sun, D. Yu, and Z. Li. 2021. Low-dose IL-2 therapy invigorates CD8+ T cells for viral control in systemic lupus erythematosus. PLoS Pathog 17:e1009858.

